# Brain network modeling with The Virtual Brain derives pharmacodynamics of ketamine

**DOI:** 10.64898/2026.02.24.707663

**Authors:** Jennifer Them, Lion Deger, Halgurd Taher, Jan Stasinski, Leon K. Martin, Jil M. Meier, Leon Stefanovski, Petra Ritter

## Abstract

Ketamine, an N-Methyl-D-aspartate receptor (NMDAR) antagonist, is used clinically as an anesthetic and antidepressant, and is also known for its psychotomimetic effects. Its impact on brain dynamics and behavior varies significantly with dosage likely via a dose-dependent modulation of the NMDARergic transmission. Currently, it is unclear how molecular changes at the microscopic level of NMDAR antagonism lead to large-scale changes in brain dynamics. We implement a dose-dependent NMDAR antagonism based on ketamine’s disinhibition theory into a biophysically grounded mean-field model within The Virtual Brain (TVB) framework to replicate ketamine’s key signatures across its dose spectrum. Our results imply that in low doses ketamine preferentially impairs excito-inhibitory neurotransmission while in higher doses antagonism on excito-excitatory connections plays a role. These findings highlight the utility of computational modeling for disentangling dose-specific mechanisms of action and provide a framework for exploring NMDAR-related interventions.

**Author summary:** Ketamine is a dissociative anesthetic at high doses, but at lower, sub-anesthetic doses, it has garnered significant interest for its rapid-acting antidepressant and anxiolytic effects. Despite its growing clinical use in psychiatric conditions, the precise neural mechanisms underlying ketamine’s dose-dependent effects remain incompletely understood. Ketamine primarily acts as a non-competitive antagonist of the NMDAR, which is expressed on both excitatory and inhibitory neurons throughout the cortex. One of the leading hypotheses explaining its antidepressant effects is the disinhibition theory which proposes that low doses of ketamine preferentially block NMDARs on inhibitory interneurons, resulting in increased cortical excitability. At high doses ketamine exerts anesthetic effects potentially through more widespread NMDAR antagonism including on excitatory neurons. In this study, we used a computational model to explore how selective NMDAR antagonism at different doses affects large-scale brain dynamics. A key novelty of our work is the integration of ketamine’s full dose spectrum within a single computational modeling framework, allowing us to relate distinct neural effects from disinhibition to anesthesia to experimental findings. This modeling approach contributes to a deeper understanding of how ketamine modulates cortical activity across different contexts.

## Introduction

Ketamine has a long history as an anesthetic drug as it modulates the glutamatergic system by acting as a N-Methyl-D-aspartate receptor (NMDAR) antagonist. Afar from its crucial role in anesthesia and emergency medicine lower, subanesthetic doses have recently been used to address treatment-resistant depression and are being investigated for a multitude of mental health disorders [1–3]. In low doses, ketamine induces a psychological state of dissociation which can be used for modeling psychosis, while in high doses it induces anesthesia. These distinct behavioral effects come with widespread changes in glutamatergic neurotransmission and large-scale electrophysiological activity [4, 5]. However, the relationship between ketamine’s dose-dependent effects and these large-scale brain dynamic changes remains unclear.

In the following, we will provide an overview of ketamine’s empirically measured effects. During ketamine anesthesia *α*-band activity decreases and increased slow-wave activity is observed in the *θ*-band (4 − 8 Hz) and *δ*-band (0.5 − 4 Hz) [6, 7]. Low doses of ketamine on the other hand show a decrease in *α*-band activity (8 − 12 Hz) and an increase in *γ*-band activity (≥ 30 Hz) [5, 8]. Anesthetic doses of ketamine acutely reduce pyramidal cell (PC) firing and glutamatergic neurotransmission in-vitro and in rodents [4, 9]. Counterintuitively, low doses of ketamine have an overall excitatory effect in rodents and humans even though ketamine antagonizes the excitatory NMDAR [10–12]. This apparent contradiction can be resolved by means of the disinhibition theory as follows [13]. On a synaptic level, ketamine impairs NMDARergic transmission but NMDARs sit abundantly on glutamatergic neurons and inhibitory interneurons (IIN) [14]. Consequently, NMDARs can impact both excitatory and inhibitory neurotransmission. In low doses ketamine seems to preferentially antagonize NMDARs located on IIN [10, 14]. This phenomenon may results from ketamine’s interaction with the NMDAR. NMDARs are blocked by Mg^2+^ which needs to be removed by predepolarization before the receptor can be opened. The binding of ketamine occurs deep within the NMDAR therefore ketamine has no effects on closed NMDARs [15]. IIN exhibit a tonic baseline activity that facilitates the removal of the Mg^2+^ block and opening of the receptor therefore ketamine can bind more easily [16–18]. Additionally, the specific subunits of NMDARs on IIN tend to have higher affinity for ketamine and lower sensitivity to their Mg^2+^ block [19, 20]. Hence, ketamine in lower doses preferentially impairs excito-inhibitory neurotransmission on IIN leading to less inhibition and therefore an increase in global excitation [14].

Izumi et al. have shown that in lower doses only a small percentage of NDMARs is blocked by ketamine, while in higher doses a large fraction of NMDARs is blocked, ensued by a complete block of NMDARs in very high doses [15]. Overall, various studies imply that low dose ketamine has opposing effects on PCs and IINs by impairing excito-inhibitory neurotransmission in the IIN leading to increased excitation in PCs and glutamate release in the cortex [10, 12, 21, 22]. In higher doses we hypothesize that ketamine can increasingly antagonize NMDARs on PCs that are open due to the prior disinhibition and increased excitation. Higher neural activity leads to a weaker Mg^2+^ block enabling modulation by ketamine. Consequently, antagonism of the NMDARs on PCs reduces excitation. These intricacies within the ketamine-NMDAR interaction necessitate to model ketamine’s effects on NMDARs on PCs and IIN distinctly depending on the dose.

Computational models of ketamine’s NMDAR antagonism are sparse but a recent study by Sacha et al. (2025) presents a mean-field model framework capturing the different mechanisms of anesthetics including ketamine and reproducing empirically observed changes seen in anesthesia [23]. Other computational studies focus either on the effects of the cortico-thalamic circuit or on the neuron level [24, 25]. No study has yet considered modeling various doses ranging from low to anesthetic within one model. Here, we use whole-brain modeling with The Virtual Brain (TVB) to identify the causal mechanisms of ketamine’s effects through an implementation of the disinhibition theory in low doses and increasing involvement of NMDARs on PCs in high doses. The brain network model (BNM) is based on the human connectome and incorporates biologically interpretable parameters. For this, brain regions are modeled by abstracted cortical columns using the Jansen-Rit (JR) neural mass model [26]. It consists of two excitatory and one inhibitory subpopulations that are coupled locally. The dynamics of a single JR model are predominately characterized by either damped or sustained oscillations. These behaviors can be described by means of bifurcation theory, therein they are associated with so-called stable foci and limit cycles (LCs). These produce *α*-band oscillations in large parts of the parameter space, while slower *θ*-band oscillations can be found in narrow regimes.

Neuromodulation induced by ketamine is modeled as an alteration of the local coupling parameters (LCPs). In detail, the disinhibition hypothesis was implemented by using sigmoidal transfer functions that translate the ketamine dose to the individual decrease in local coupling between the neuron types. As a target to validate the model we choose to combine effects found on the electrophysiological and subcellular level. Specifically, we investigated changes in *α*- and *θ*-band power of post-synaptic potentials (PSPs) as well as the firing rate of the PCs, which acts as a proxy for excitation. To our knowledge this is the first study that incorporates ketamine’s dose-dependent mechanisms of action in a whole-brain model. Moreover, it allows to perform an in-depth analysis of the local dynamics as well as network effects, which are exhibited at different doses of ketamine using numerical bifurcation analysis.

The organization of the paper is as follows. First, we study how independent changes of local coupling between subpopulations impact the targets given by the PSP band powers and the firing rate. Second, we perform an analysis of the effects of uniform NMDAR-antagonism, meaning the LCPs are changed in the same linear manner by ketamine. We show that electrophysiological effects but not changes in firing rate of the PCs can be produced by this uniform NMDAR antagonism. Third, we implement the preferential antagonism of NMDARs on IIN in low doses and the progressive involvement of the NMDAR antagonism on excito-excitatory LCPs with increasing ketamine dose. This is realized through individual transfer functions to model ketamine’s impact on the decrease in NMDARergic LCPs. With this approach we reproduced the changes in synaptic and electrophysiological properties in line with literature. Lastly, we conclude what this work can contribute to the understanding of ketamine’s drug action to improve future treatments.

## Materials and methods

### Data

For our network simulation we used the s900 release of the structural connectivity data from the Human Connectome Project (HCP), a publicly available dataset of 785 young adults [27]. The downloaded dataset was preprocessed with the minimal preprocessing pipeline for HCP [28]. The segmentation was done in FreeSurfer, the cortical regions of interest were defined using the Desikan-Killiany parcellation [29], while subcortical regions were delineated with the Fischl atlas [30], resulting in a total of 82 regions of interest. Diffusion tensor imaging (DTI) is used to obtain a structural connectome from diffusion-weighted MRI consisting of a weight matrix *W* and a tract length matrix *TL* that were averaged across subjects. *W* is given by the entries *W*_*kl*_ that denote the fiber counts between regions of interest (ROIs) *k* and *l*. For simulation the weights are log-transformed and normalized as done in Eq. (1), to obtain the coupling matrix *C*.

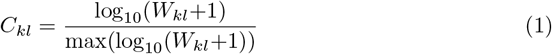

*TL* is used to calculate the time delays *τ*_*kl*_ between regions *k* and *l*,

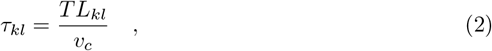

where *v*_*c*_ is the conduction speed.

### Brain network model

Using The Virtual Brain framework [31], we here employ a BNM to simulate spontaneous neural activity of each brain region. The behavior of each region is captured by the JR neural mass model [26]. A neural mass represents the average activity of a population of neurons. In the JR model a neural mass consists of one PC, one excitatory interneuron (EIN) and one inhibitory interneuron (IIN) that are locally coupled (see Fig. 1). The PC projects to both EIN and IIN and both interneuron populations project back to the PC population. The dynamics of a single JR model with the three local populations are given in Eqs. (3) to (8).

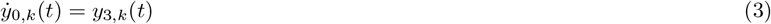

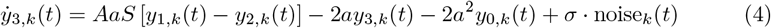

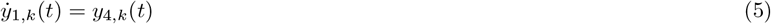

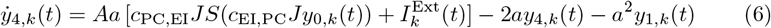

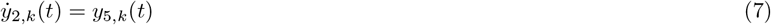

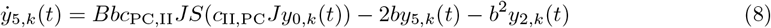

**Fig 1.**
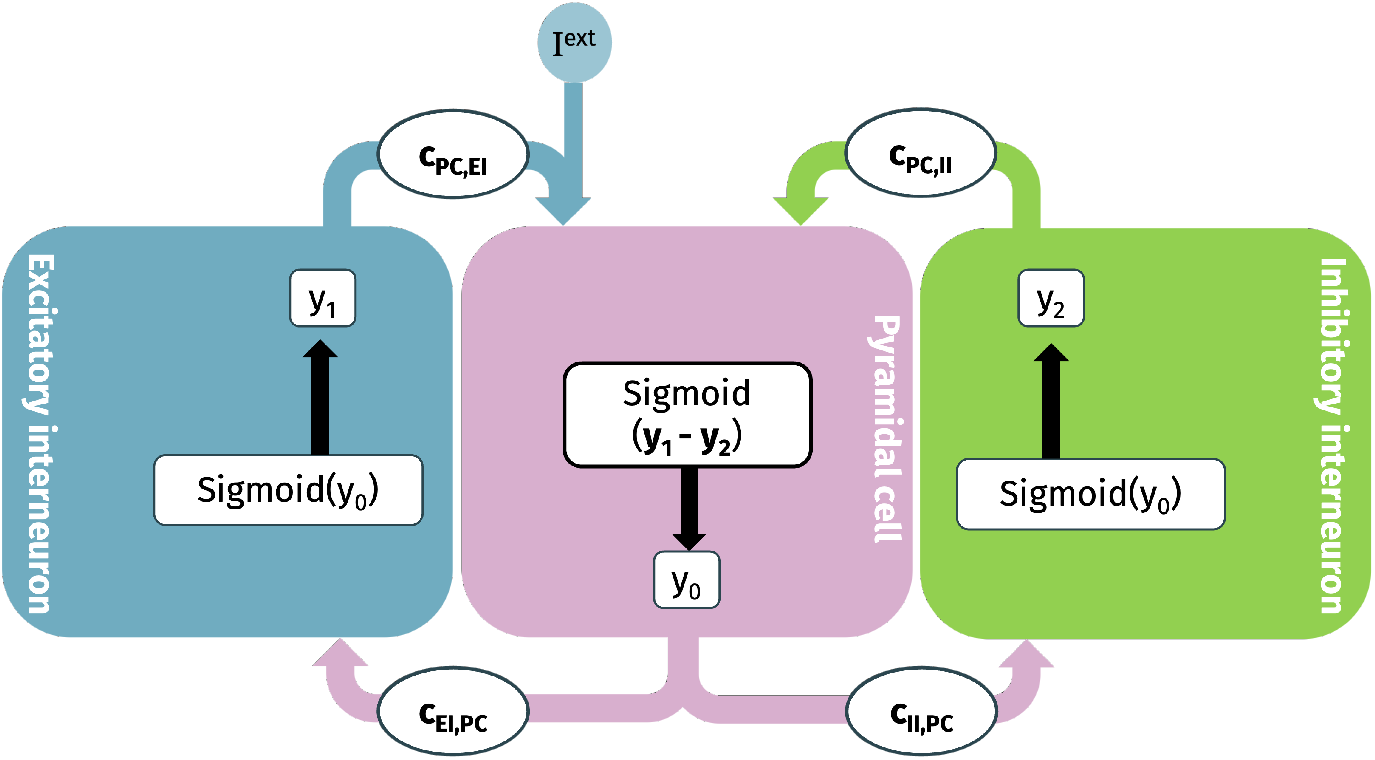
Schematic representation of the JR neural model. The JR model consists of three subpopulations: PCs, EINs and IINs that are locally coupled by *c*_EI,PC_, *c*_PC,EI_, *c*_II,PC_ and *c*_PC,II_. Potential is converted to firing rate in each subpopulation by a sigmoidal function *S*(*v*) [26]

Here, *y*_0,*k*_ and *y*_3,*k*_ represent the PC, *y*_1,*k*_ and *y*_4,*k*_ the EIN and *y*_2_, *k* and *y*_5,_ the IIN sub-populations. The region index is denoted by *k* = 1, …, *N*, where *N* is the number ROIs. The parameters *A* and *B* are the maximum excitatory and inhibitory postsynaptic membrane potentials, respectively, whereas *a* and *b* are the reciprocal of the excitatory and inhibitory time constants, respectively. Each region receives the independent Gaussian white noise noise_*k*_(*t*) which enters in Eq. (4) with the amplitude *σ*.

The local coupling strengths between subpopulations are denoted by *c*_EI,PC_, *c*_PC,EI_, *c*_II,PC_ and *c*_PC,II_ which reflect the number of synapses of their respective biological neuron type. Coupling is realized through a sigmoidal transfer function which translates from pre-synaptic potential *v* to post-synaptic firing rate *S*(*v*):

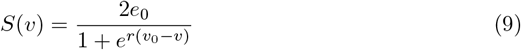

where *e*_0_ represents the maximum firing rate of the population, *v*_0_ denotes the post-synaptic potential (PSP) at which 50% of the maximum firing rate is achieved and *r* describes the slope of the sigmoidal function.

Moreover, the time-dependent external 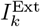 drives region *k* and is constant, when a neural mass in isolation is considered. In presence of long-range coupling among ROIs however, it includes a weighted sum of incoming firing rates which are computed via *S*(*v*), the long-range coupling matrix *C* and tract length matrix *TL*, resulting in Eq. (10).

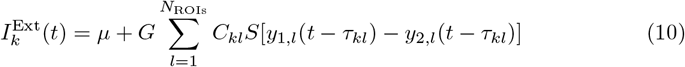

Here, *G* is used to scale the long-range connections, whereas *µ* is an external background activity common to all ROIs. All parameter values for the BNM can be found in Table 1.

**Table 1.** BNM simulation parameters, which are fixed throughout this work, if not stated differently.

| Symbol | Description | Value |
| --- | --- | --- |
| $G$ | Global coupling | 1.95 |
| $v_c$ | Conduction speed | $8 \text{ m} \cdot \text{s}^{-1}$ |
| $\mu$ | External input | $0.01 \text{ ms}^{-1}$ |
| $J$ | Synaptic weight | 135.0 |
| $\sigma$ | Noise amplitude | $10^{-8}$ |
| $A$ | Maximum amplitude excitatory PSP | 3.25 mV |
| $B$ | Maximum amplitude of inhibitory PSP | 22.0 mV |
| $a$ | Reciprocal of excitatory time constant | $0.1 \text{ ms}^{-1}$ |
| $b$ | Reciprocal of inhibitory time constant | $0.05 \text{ ms}^{-1}$ |
| $v_0$ | Firing threshold | 6.0 mV |
| $r$ | Steepness of sigmoidal | $0.56 \text{ mV}^{-1}$ |
| $c_{\text{EI,PC}}$ | Local coupling from PC to EI | 1.0 |
| $c_{\text{PC,EI}}$ | Local coupling from EI to PC | 0.8 |
| $c_{\text{II,PC}}$ | Local coupling from PC to II | 0.26 |
| $c_{\text{PC,II}}$ | Local coupling from II to PC | 0.25 |
| $e_0$ | Maximum firing rate of neural population | $0.0025 \text{ ms}^{-1}$ |

### Bifurcation analysis

To investigate the dynamical landscape that underlies the neural mass model, we perform a numerical bifurcation analysis of a single JR model in absence of long range coupling, i.e, 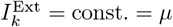, using auto-07p [32]. Therefore, we will omit the region index *k* in the following. The bifurcation diagram in Fig. 2 displays the postsynaptic potential (PSP) of the pyramidal cell given by PSP = *y*_1_ − *y*_2_ as a function of the external input *I*^Ext^. For small values of *I*^Ext^ ⪅ 0.1148 ms^−1^ there exists a stable fixed points (FP) branch. By increasing *I*^Ext^ this branch undergoes a saddle-node bifurcation (SN) that folds and destabilizes the branch. A second SN occurs with no change in stability. The unstable FP branch undergoes a Andronov-Hopf bifurcation at *I*^Ext^ ≡ *I*_hb_ ≈ 0.4024 ms^−1^ that gives rise to stable limit cycles (LCs) with frequencies in the *α*-band, as can be seen in Fig. 2B. The stable LC branch undergoes a SN bifurcation of LCs at 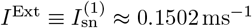 and 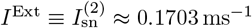, such that the branch first destabilizes and then stabilizes again. Overall, two families of stable LCs coexist in the range 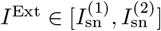 exhibiting *θ*- and *α*-band activity, respectively (see Fig. 2B). For the BNM simulation, we set the constant external input to *µ* = 0.01ms^−1^ such that a node in absence of long-range coupling would possess only one stable fixed point. Nevertheless more complex dynamics can be induced in the BNM, since long range coupling will shift the individual regions towards larger *I*^Ext^, as we show in Section Sigmoidal implementation of ketamine is crucial to reproduce low-dose and anesthetic effects.

**Fig 2.**
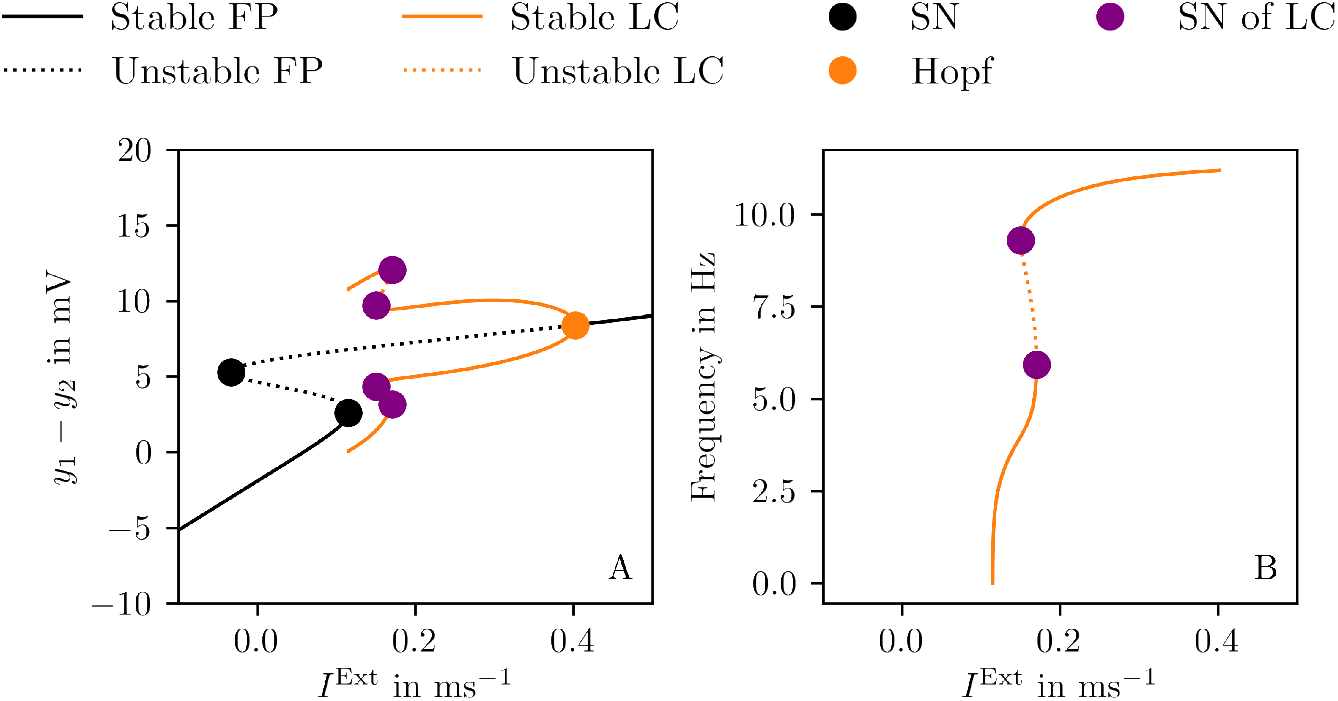
Bifurcation diagram of the Jansen-Rit neural mass model. **(A)**Postsynaptic potential PSP = *y*_1_ −*y*_2_ of JR vs. *I*_Ext_ for fixed points and limit cycles (LC) and (**B**) frequencies of LC vs. *I*^Ext^. Solid lines display stable branches while dashed lines display unstable branches. Black lines show fixed points (FPs) with black dots indicating saddle-node (SN) bifurcations, orange lines show minimum and maximum of LCs with orange dots indicating Andronov-Hopf bifurcations. Purple dots show saddle-node bifurcation of the LCs. Parameter values can be found in table 1. Bifurcation analysis was done with the software auto-07p [32].

### Transformation of local coupling parameters

We implement ketamine’s NMDAR-antagonism as an alteration of the LCPs *c*_EI,PC_, *c*_PC,EI_ and *c*_II,PC_. To understand the effects of LCP variations on the firing rate and spectral measures, a parameter space exploration was performed for *c*_EI,PC_ ∈ [0.5, 1.5], *c*_PC,EI_ ∈ [0.4, 1.2] and *c*_II,PC_ ∈ [0.125, 0.375] using an equidistant 20 × 20 × 20 grid. Two different trajectories *c*_*ij*_(Ket) as a function of ketamine dose Ket were introduced in this parameter space, with (*i, j*) ∈ {(PC, EI), (EI, PC), (II, PC)}.

### Linear transfer function

We implemented the linear trajectory 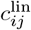 by defining a uniform scaling factor (1-Ket) that scales the default values 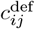of the LCPs. We will refer to this scaling defined in Eq. (11) as uniform scaling.

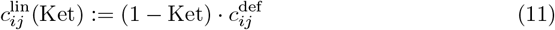

Here, 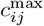 is the maximum of each LCP at Ket=0.0. The corresponding intervals of LCPs are *c*_EI,PC_ ∈ [0.0, 1.00], *c*_PC,EI_ ∈ [0.0, 0.800] and *c*_II,PC_ ∈ [0.0, 0.2571] (see also Fig. 6).

### Sigmoidal transfer function

Additionally, we implemented a sigmoidal trajectory 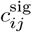by constructing inverted sigmoidal functions with distinct parameter values for each LCP. We will refer to this scaling defined in Eq. (12) as sigmoidal scaling.

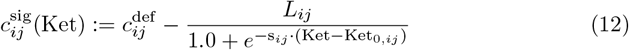

*L* is the range between maximum 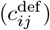 and minimum coupling value, *s*_*ij*_ controls the steepness and Ket_0,*ij*_ determines the inflection point of the curve. To account for differential sensitivity to ketamine, we set the inflection point Ket_0,*ij*_ of *c*_II,PC_ to a lower Ket value than for *c*_EI,PC_ and *c*_PC,EI_. The parameter values are provided in Table 2.

**Table 2.**
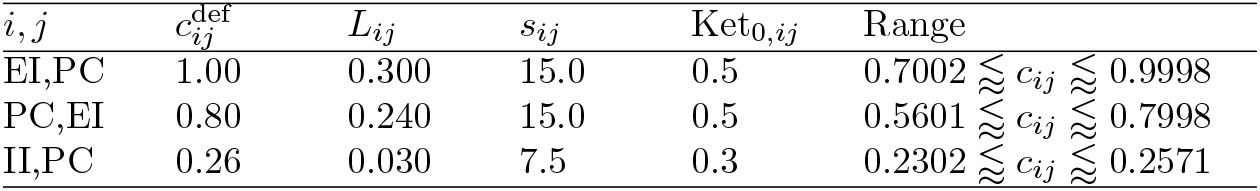
Parameter values for the sigmoidal transfer function 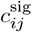. The table shows the default coupling value 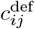, difference between maximum and minimum coupling value *L*_*ij*_, steepness of the sigmoid *s*_*ij*_ and inflection point of the curve Ket_0,*ij*_ for each LCP. These parameters define the sigmoidal functions used to model ketamine-induced changes in local coupling. Note that minimum and maximum values are never reached due to the asymptotic nature of the sigmoid

### Simulation

The JR model is a nonlinear system, with a single JR node exhibiting multistability (as detailed in Bifurcation analysis). When extended to a BNM, the system becomes high-dimensional, further increasing its complexity. Additionally, the simulations are stochastic such that different noise realizations need to be explored. Overall, to ensure robustness of the results and capture variability, ten trials with varying initial conditions and noise realizations were conducted. The initial conditions were drawn uniformly random within the ranges of *y*_0_ ∈ [0.09, 0.11], *y*_1_ ∈ [0.0, 23.0], *y*_2_ ∈ [0.0, 16.0], *y*_3_ ∈ [ − 0.002, 0.002], *y*_4_ ∈ [ − 0.04, 0.04], *y*_5_ ∈ [ − 0.1, 0.1]. Each simulation was run for 120 s using the Heun integration scheme and an integration time step of *dt* = 1 ms. A transient of 115 s was cut-off leaving 5 s of the simulated neural signal for concurrent analysis steps. We explored 1000 equidistant ketamine values given Ket ∈ [0.0, 1.0].

### Spectral properties of simulated neural signal

To analyze the spectral properties of the simulated signals, we examine their frequency composition and power distribution. We start by extracting the PSP of the PC of each region *k* by subtracting the PSP_*k*_ of the IIN *y*_2_ from the PSP_*k*_ of the EIN *y*_1_ (PSP_*k*_ = *y*_1,*k*_ − *y*_2,*k*_). Then, we calculate the power spectral densities (PSDs) from the PSP_*k*_ using the Fourier transform. The resulting PSD_*k*_(*f*) = ℱ (PSP_*k*_)(*f*) are averaged across all regions to characterize the spectral properties of the entire system, resulting in 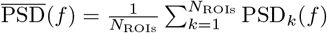. To quantify power within key frequency bands, we computed the power within the *α*-band (8 Hz *< f* ≤ 12 Hz) and the *θ*-band (4 Hz < *f* ≤ 8 Hz) by integrating 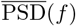 over the corresponding ranges.

## Results

In this study we developed a BNM to understand ketamine’s main mechanism of action, namely the antagonism of the NMDAR. The BNM comprises a standardized human SC in which each brain region is modeled by a neural mass. Each neural mass represents a cortical column with three sub-populations, namely PCs, EINs and IINs, that are interconnected by the LCPs. Here, ketamine’s NMDAR antagonism is implemented as an alteration of these LCPs. They will be altered as functions *c*_*i,j*_(Ket) of the ketamine dose Ket, where *c*_*i,j*_(Ket) can be either linear or sigmoidal, as described in Section Transformation of local coupling parameters. The JR neural mass model is a high-dimensional, non-linear model with a complex bifurcation landscape as described in Section Bifurcation analysis. The bifurcation diagram is given as a function of *I*^Ext^, but under the assumption of fixed LCPs. While periodic oscillations in the *α*- and *θ*-band can be expected for default LCP values, it is unclear how the system will behave when they are varied. In the following we will provide an analysis of the spectral properties and average firing rate of the BNM, empirically studied measures of ketamine’s dose-dependent effects, under variations of the relevant local parameters *c*_EI,PC_, *c*_PC,EI_ and *c*_II,PC_.

### Independent change of local coupling reveals distinct spectral areas

It is essential to initially consider each parameter independently to assess the impact on firing rate and spectral properties of the BNM. We investigate the firing rate present at each point of the parameter grid, as shown in Fig. 3. Here, we find that increasing *c*_PC,EI_ shifts the window of high mean firing rate along the axes of *c*_EI,PC_ and *c*_II,PC_. When *c*_PC,EI_ is fixed, the mean firing rate increases with increasing *c*_EI,PC_ and decreasing *c*_II,PC_. As an example, for a value *c*_PC,EI_ = 0.40 of the excito-excitatory coupling, we find a narrow region of the other LCPs, *c*_EI,PC_ ≥ 0.82, *c*_II,PC_ ≤ 0.19, for which the firing rate reaches up to 4.6 Hz. Increasing *c*_PC,EI_ shifts the region’s boundary towards lower *c*_EI,PC_ and higher *c*_II,PC_, thereby increases its size and the maximum firing. *c*_PC,EI_ and *c*_EI,PC_ are excito-excitatory coupling parameters, increasing them will lead to higher excitatory activity. This can be compensated by more inhibition, hence by increasing the excito-inhibitory coupling *c*_II,PC_.

**Fig 3.**
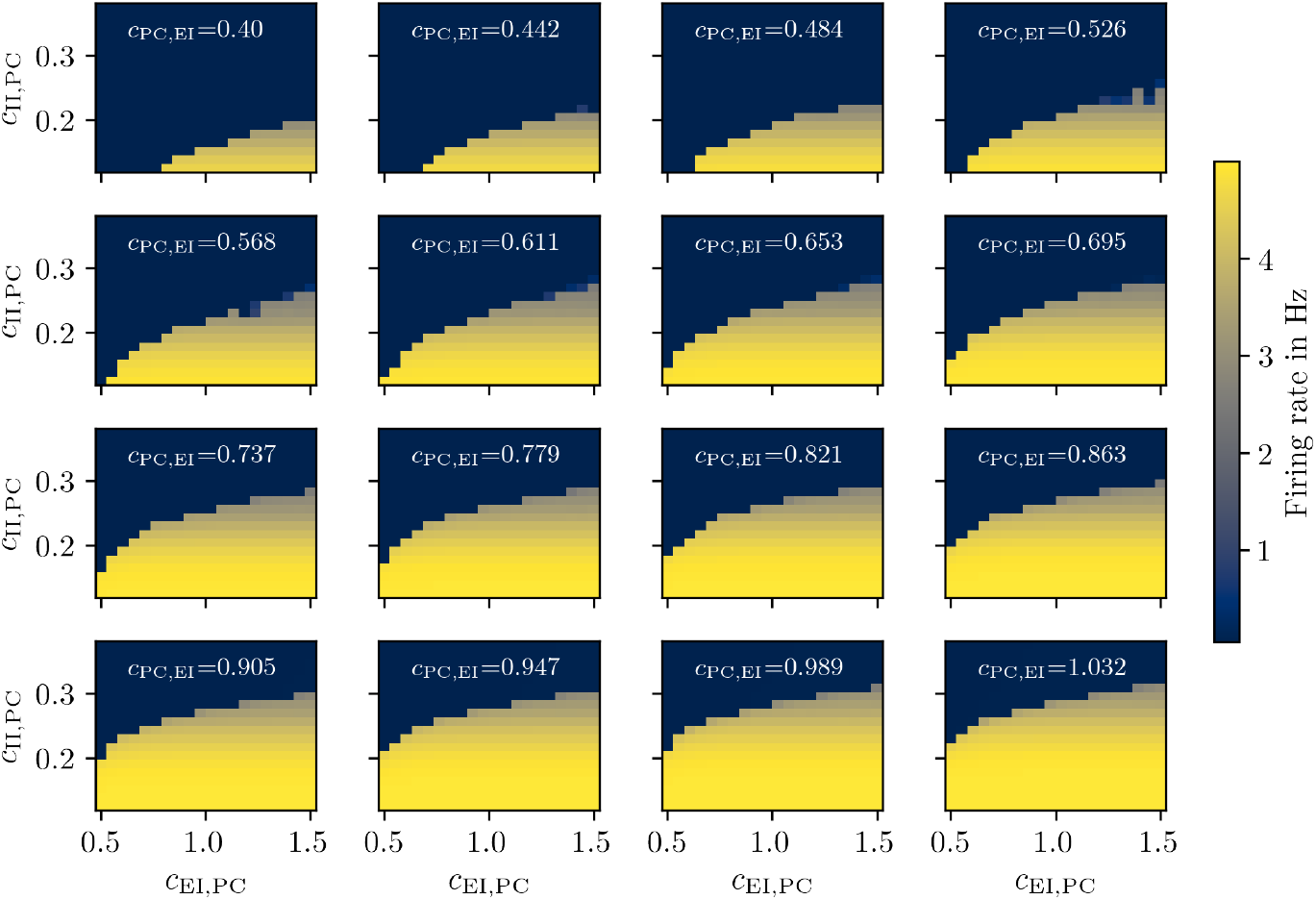
Firing rate of the pyramidal cells for different combinations of LCPs. The mean firing rate is computed over all regions and color coded for different combinations of the parameters *c*_EI,PC_, *c*_PC,EI_, *c*_II,PC_. Each panel shows a cross section in the (*c*_EI,PC_, *c*_II,PC_)-plane given different values of *c*_PC,EI_. In each panel a critical boundary is present marking an abrupt decrease in firing rate.

The *α*-band power is depicted in Fig. 4. While at *c*_PC,EI_ = 0.40 regions of relevant alpha power are absent, larger excito-excitatory coupling induces oscillations in this band. As an example, at *c*_PC,EI_ = 0.779 we find that the *α*-power reaches up to 1.59 mV^2^/Hz in a triangular region delimited by *c*_EI,PC_ ≥ 0.974 and 0.257 ≤ c_II,PC_ ≤ 0.283. Similar to the firing rate, increasing *c*_PC,EI_ increases the size of this region as well as the maximum *α* power reached.

**Fig 4.**
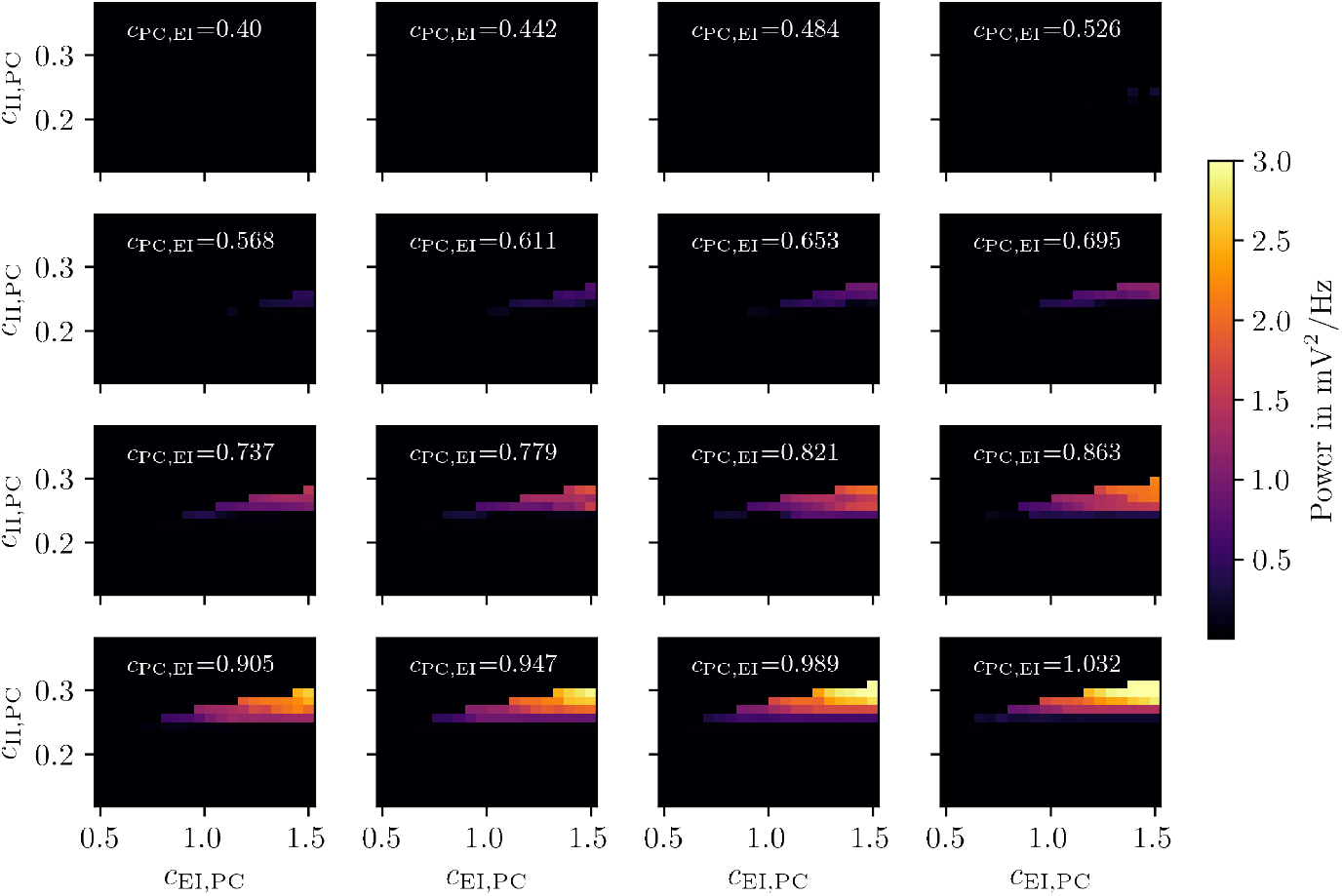
*α*-band power for different combinations of LCPs. The mean *α*-band (8-12 Hz) power is computed across all regions and color coded for different combinations of the parameters *c*_EI,PC_, *c*_PC,EI_, *c*_II,PC_. Each panel shows a cross-section in the (*c*_EI,PC_, *c*_II,PC_) plane given different values of (*c*_PC,EI_).

Finally, the *θ*-band power is shown in Fig. 5. Analogous to the *α*-band, we find increased theta power for larger *c*_PC,EI_, e.g., *c*_PC,EI_ = 0.737 with the *θ*-power reaching up to 1.44 mV^2^/Hz. In contrast to the firing rate and *α*-band power however, the *θ*-band power does not monotonously increase when increasing *c*_PC,EI_. Instead, it peaks at *c*_PC,EI_ = 0.737 before decreasing. Similarly, the size of the *θ*-power region increases up to this point and then declines.

**Fig 5.**
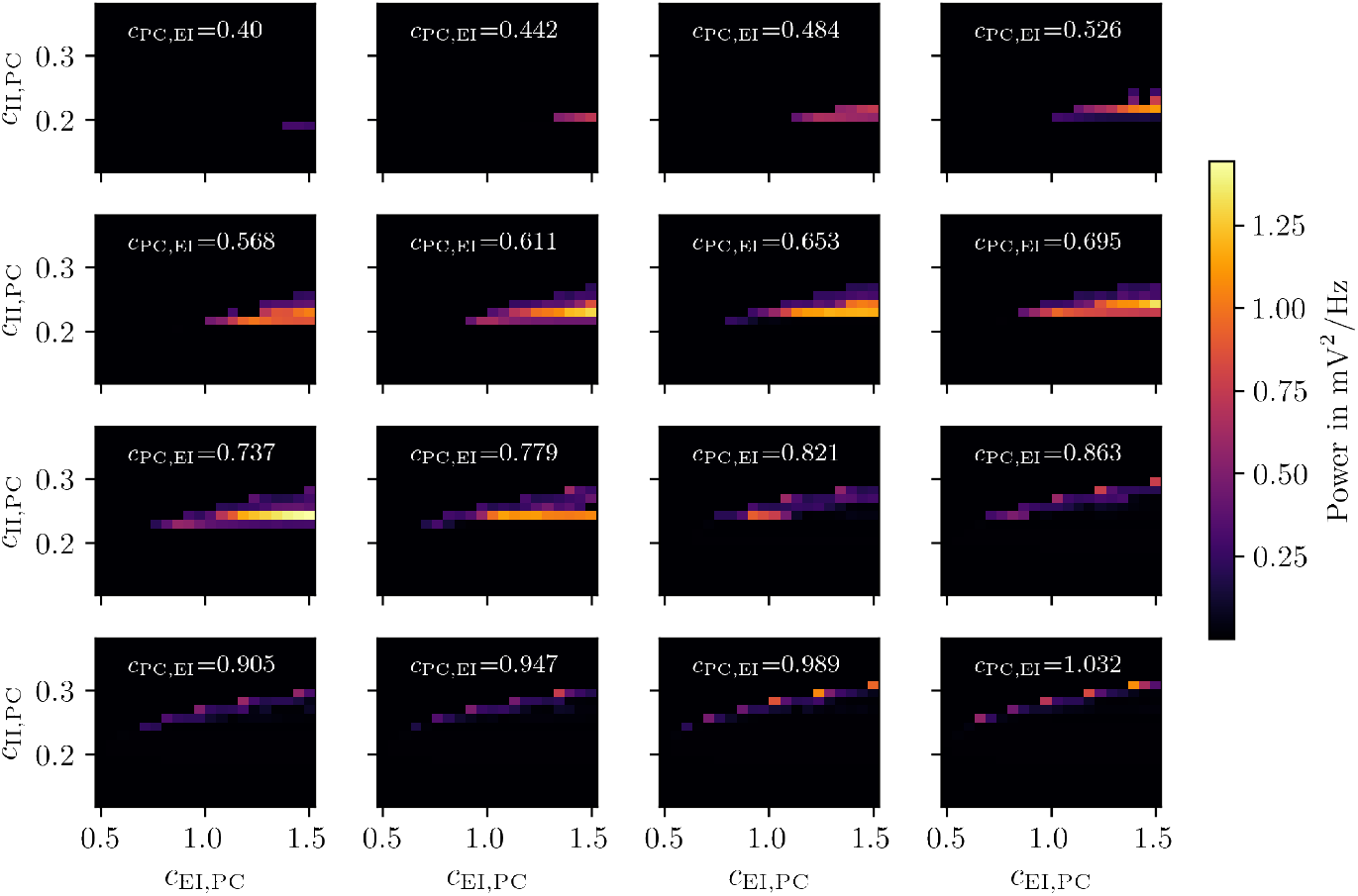
*θ*-band power for different combinations of local coupling parameters. The mean *θ* power (4-8 Hz) is computed over all regions and color coded for different combinations of the parameters *c*_EI,PC_, *c*_PC,EI_, *c*_II,PC_. Each panel shows a cross-section in the (*c*_EI,PC_, *c*_II,PC_) plane given different values of (*c*_PC,EI_).

In summary, the average firing rate, as well as *α*- and *θ*-band powers are highly sensitive to LCP variations, revealing distinct regimes with varying firing rates and non-overlapping band power distributions. This exploration allows us to identify the lower and upper bounds of LCPs where changes in firing rate and spectral properties emerge. Notably, relevant *α*- and *θ*-power appear within distinct parameter ranges.

Based on this parameter exploration of the coupled JR model, we introduce theories about ketamine’s drug mechanisms. For this we define different parameter space trajectories (*c*_EI,PC_(Ket), *c*_PC,EI_(Ket), *c*_II,PC_)(Ket) as described in Transformation of local coupling parameters.

### Linear scaling of NMDARergic coupling fails to produce changes in firing rate

We start with the assumption that ketamine antagonizes all NMDARs uniformly, i.e., the trajectory is linear. The dependency of 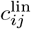 on Ket is shown in Fig. 6. In Fig. 7 the impact of this uniform linear scaling on the different measures, namely firing rate, *α*-band and *θ*-band power, is depicted.

**Fig 6.**
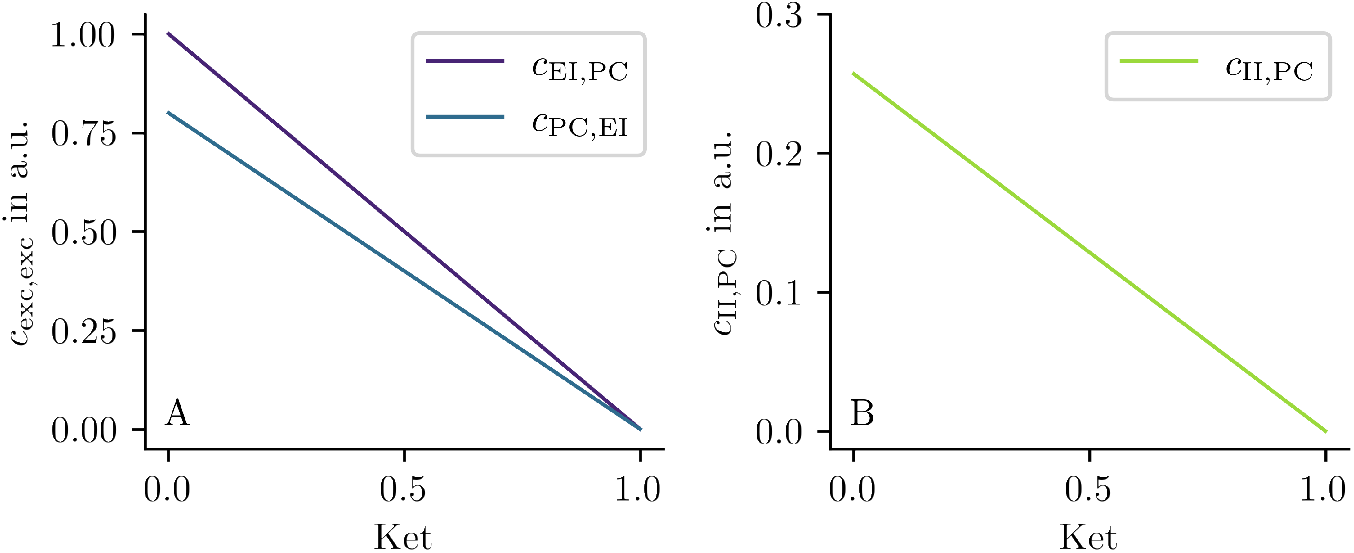
Effects of uniform NMDAR-antagonism on the LCP. Values of LCP *c*_EI,PC_, *c*_PC,EI_ (**A**) and *c*_II,PC_ (**B**) vs. ketamine parameter. Value of local coupling parameters decreases linearly with increasing ketamine parameter. The transformation is a linear function with *c*_*ij*_ = (1 − Ket) · 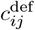 for (*i, j*) ∈ {(PC, EI), (EI, PC), (II, PC)}.

**Fig 7.**
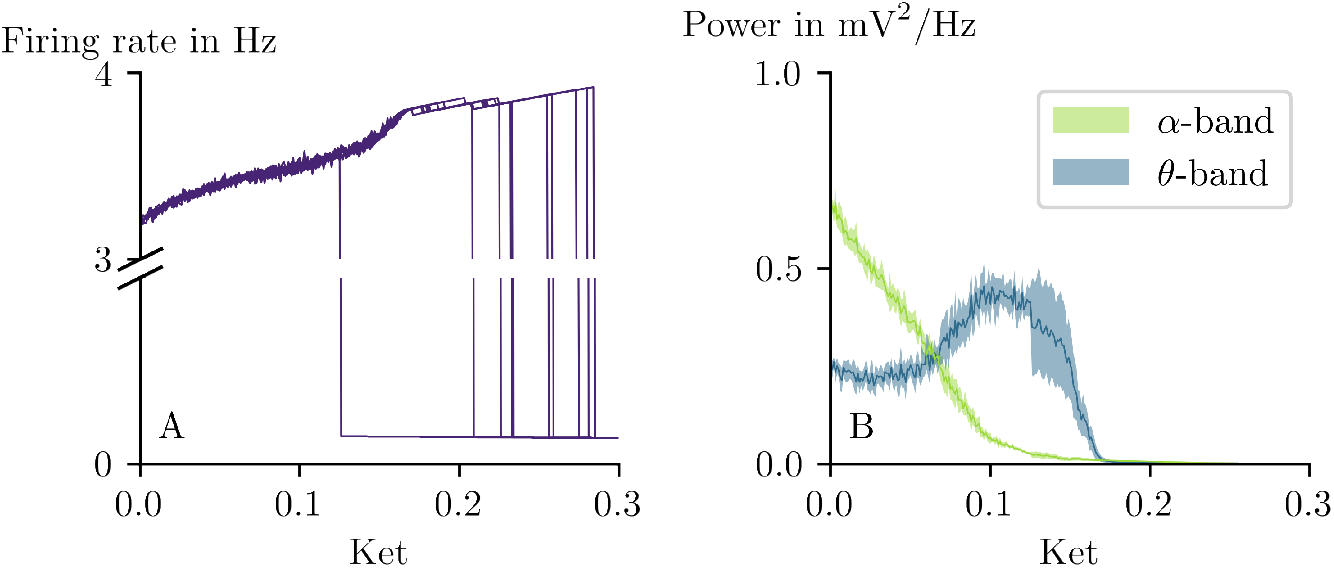
Effects of uniform NMDAR-antagonism on firing rate and spectral measures. The ketamine parameter has an uniform effect on all LCP. The mean firing rate, *α*-band power and *θ*-band power are computed over all regions for different combinations of the parameters *c*_EI,PC_, *c*_PC,EI_, *c*_II,PC_. (**A**) Firing rate is shown for ten trials vs. Ket. The firing rate increases with increasing Ket until a critical boundary is reached causing the firing rate to abruptly decrease. (**B**) Mean and standard deviation of the *α*-band (green) and *θ*-band power (blue) are shown for ten trials vs. Ket. Power in the *α*-band decreases across Ket range while power in the *θ*-band firstly increases peaking around Ket = 0.121 and then decreases with increasing Ket value. Both *α* and *θ*-band power approach zero for Ket *>* 0.2.

As can be seen in Fig. 7A, in absence of ketamine (Ket = 0.0) the BNM exhibits a mean firing rate of 3.207 Hz across regions and trials. Increasing Ket leads to an increase in firing rate until a critical point is reached and the firing rate sharply declines to nearly zero. The precise Ket value at which this transition on the network level occurs varies across trials due to the system’s multistability together with differing initial conditions and noise realizations. These factors lead to distinct network states, causing transitions to occur at different ketamine doses.

Regarding the spectral properties (Fig. 7B), we observe that in the absence of ketamine (Ket = 0.0), the mean *α*-power across trials is 0.51 mV^2^/Hz, while the mean *θ*-power is 0.25 mV^2^/Hz. As Ket increases, the mean *α*-power progressively declines, whereas the mean *θ*-power initially shows a slight decrease before increasing to a peak of 0.35 mV^2^/Hz. This peak occurs when the mean *α*-power is low (0.05 mV^2^/Hz). Beyond this point, the mean *θ*-power begins to decline.

Notably, the uniform antagonism induces slowing from the *α*- to the *θ*-band similar to what was reported in literature [6, 7]. However, in the regions of elevated firing rates, the *α*- and *θ*-band powers are offset in parameter space with respect to each other and have different extents. Hence, when increasing Ket, changes of these measures do not occur simultaneously, but onset at distinct Ket values. There is already a level of selectivity and dose-dependency of the output measures but realistic changes in firing rate are missing.

### Sigmoidal implementation of ketamine is crucial to reproduce low-dose and anesthetic effects

The uniform linear scaling was an intuitive approach of the LCP alterations. Here, we want to consider hypothesized biological mechanisms of action more in detail, namely the disinhibition hypothesis. NMDARs on different neuron types have distinct properties impacting their interaction with ketamine including the strength of the Mg^2+^ block and affinity for the ketamine molecule. We capture this dose-dependent, selective NDMAR antagonism by defining a more complex *c*_*ij*_ vs. Ket relationship as follows. In line with the disinhibition hypothesis, lower values of Ket should have higher impact on the decrease of *c*_II,PC_. Larger values on the other hand should lead to the onset of changes in the other two LCPs *c*_EI,PC_ and *c*_PC,EI_. This behavior of the NMDARergic LCP can be realized by distinct sigmoidal transfer functions as described in Eq. (12). The sigmoids allow to control the Ket values at which the effects come into play, as depicted in Fig. 8. The local coupling from pyramidal cells to inhibitory interneurons, *c*_II,PC_, starts slightly below the default value of *c*_II,PC_ = 0.26 in absence of ketamine, namely 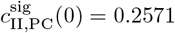. It decreases quickly until the inflection point at Ket_0_ = 0.3 and then asymptotically approaches a minimum value (Fig. 8B). For more details see Section Transformation of local coupling parameters). The other excito-exitatory LCPs *c*_EI,PC_ and *c*_PC,EI_ shown in Fig. 8A remain nearby their maximum values 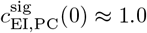 and 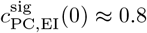 for low Ket and decrease rapidly near Ket ≊ 0.5.

**Fig 8.**
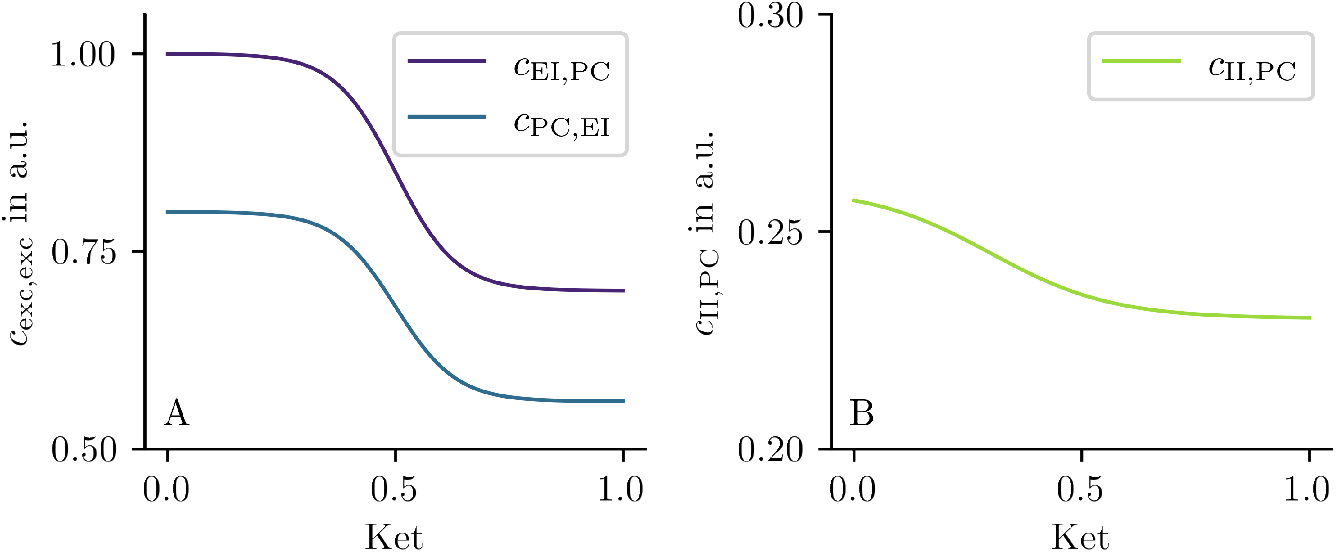
Changes in NMDARergic coupling parameters induced by ketamine. These sigmoidal functions describe the value of the LCPs *c*_EI,PC_, *c*_PC,EI_ (**A**) and *c*_II,PC_ as a function of the ketamine parameter. The main difference between excito-exitatory (*c*_EI,PC_, *c*_PC,EI_) and excito-inhibitory *c*_II,PC_ LCP is the difference of inflection points of the sigmoid. Detailed values are given in Table 2.

The impact of using selective sigmoidal instead of identical linear transfer functions on firing rate and *α*- and *θ*-band powers is shown in Fig. 9A and B, respectively. We observe that low ketamine values initially lead to an increase in firing rate peaking at a mean firing rate across trials of *r*_max_ = 3.527 Hz at Ket = Ket_max_ = 0.329. For larger Ket values the firing rate decreases (see Fig. 9A). Additionally, we observe a decrease in *α*-band power for low Ket values and an increase in *θ*-band power (see Fig. 9B) as the Ket dose increases. *θ*-band power peaks at 0.49 mV^2^/Hz for Ket = 0.361. The Ketamine value at which the mean *θ*-power surpasses the mean *α*-power is nearby Ket_max_. Given that ketamine blocks an excitatory receptor the increase in firing rate in low doses seems counterintuitive. However, it is reasonable to assume that Ketamine due to its site-dependent effects alters the local excitation–inhibition (E/I) ratio in a dose-dependent manner. Indeed, as shown in panels C and D of Fig. 9, the E/I ratio increases initially until Ket ≈ Ket_max_ before decreasing. This is a direct consequence of the implemented disinhibition hypothesis and results in the behavior of the firing rate observed for increasingly higher doses of Ketamine.

**Fig 9.**
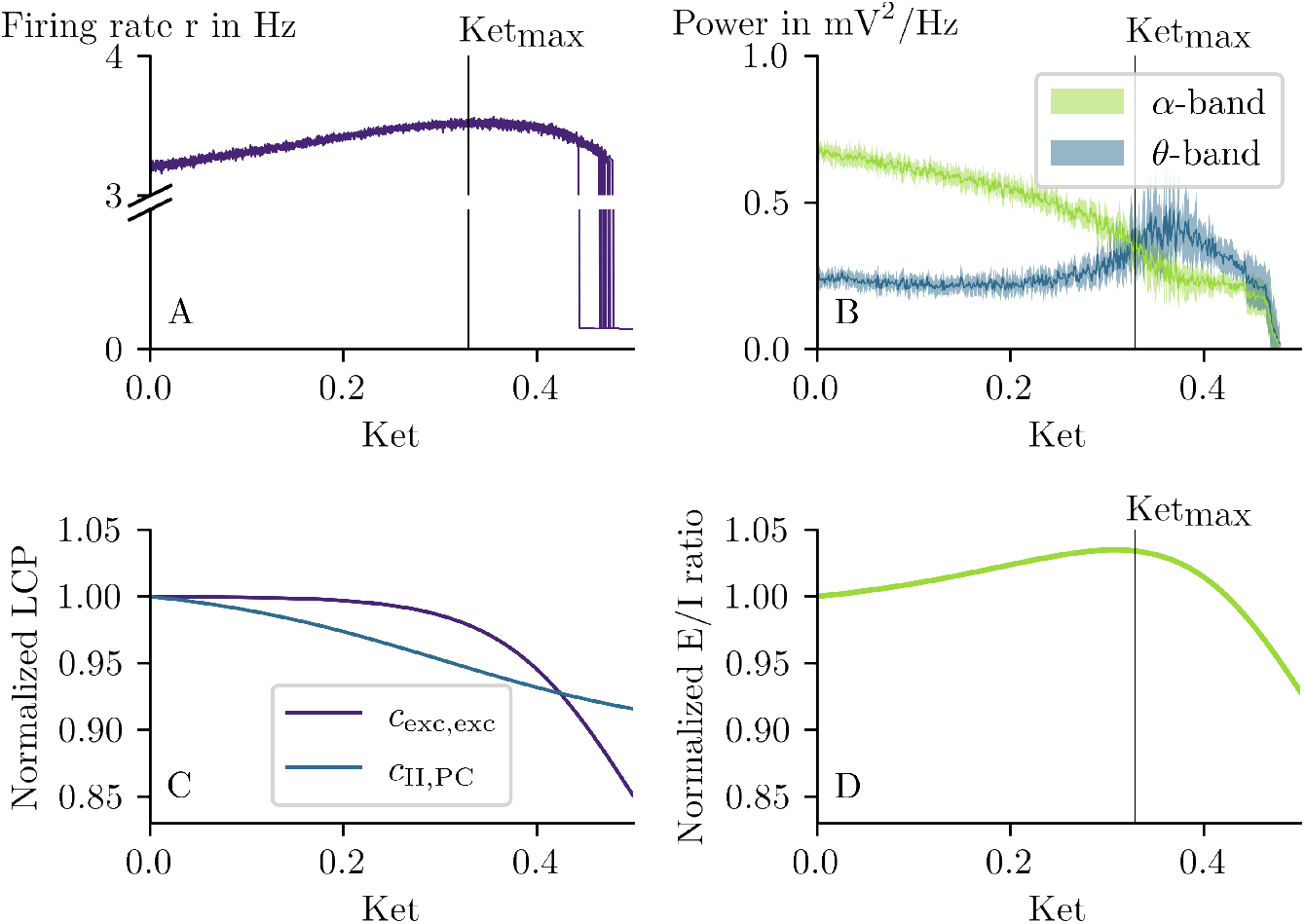
Effects of selective NMDAR-antagonism on the network. (**A**) Selective NMDAR-antagonism firstly produces an increase followed by a slow decrease in firing rate by increasing Ket. Firing rate is shown for ten trials as a function of the Ket. The maximum firing rate across ten trials *r*_max_ = 3.527 Hz is reached at Ket_max_ = 0.329. (**B**) Power in *α*- and *θ*-band is shown. The *α*-band power decreases while the *θ*-band power increases peaking at Ket = 0.361 and then decreases again with increasing ketamine dose Ket. In (**C**) the normalized LCP values and in (**D**) the respective normalized E/I ratio are shown. By construction, *c*_II,PC_ declines faster than the excito-excitatory LCP. This leads to an initial increase of the local E/I ratio that peaks at Ket = 0.308, before falling off.

Compared to the uniform scaling 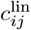, the sigmoidal scaling 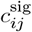introduces additional selectivity, aligning with the disinhibition hypothesis by differentially modulating excito-excitatory and exctio-inhibitory connections. This distinction is crucial, as 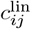 fails to reproduce the empirically observed changes in firing rate, despite already exhibiting a degree of selectivity and a dose-dependent effect on output measures (as shown in Fig. 7). In contrast, 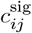 successfully captures the dose-dependent empirical changes, both in firing rate [4] and in spectral properties [6, 7]. Next, we further analyzed the effects of Ket at the single-node level providing a clearer overview of its effects without the impact of network interactions.

The bifurcation diagram provided in Fig. 2 in Section Bifurcation analysis is for Ket = 0.0, for which the LCPs are given by their default values. Altering Ket in a single node leads to changes of the bifurcation structure that underpins the network dynamics. To understand the impact of the Ket parameter on the dynamics, single JR PSP vs. *I*^Ext^ bifurcation diagrams are computed for selective Ket values in the presence of 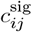, as shown in Fig. 10.

**Fig 10.**
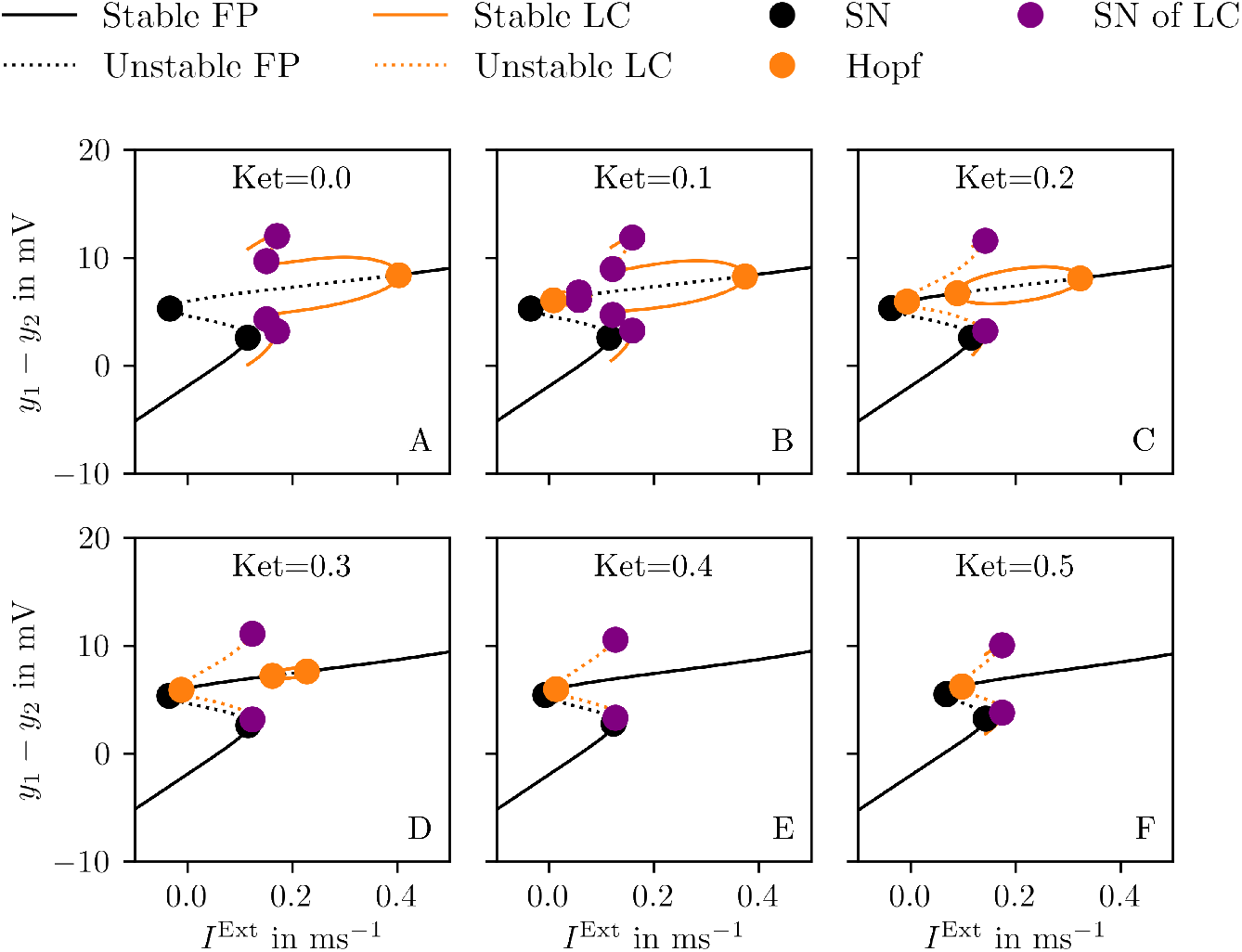
Single node bifurcation diagrams for varying doses of ketamine. Bifurcation diagrams showing PSP = *y*_1_− *y*_2_ as a function of constant external input *I*^Ext^ for different values of Ket in a range of 0.0 to 0.5 in steps of 0.1. Increasing Ket decreases the range of *I*^Ext^ values for which the LCs associated with *α*-band oscillations can be found. At the same time, it shrinks the stable part of the *θ*-band LC. Colors and symbols are as described in Fig. 2.

In absence of ketamine (Ket = 0.0, Fig. 10A), we find a Hopf bifurcation that gives rise to fast LCs (*α*-band) and two consecutive saddle-node bifurcations of LCs that give rise to the stable branch of slow LCs (*θ*-band). Here, the unstable foci situated between the SN at *I*^Ext^ ≈ −0.0333 ms^−1^ and Hopf bifurcation at *I*^Ext^ ≈ 0.4024 ms^−1^, become stable across an increasing range of *I*^Ext^ as the branch of the fast LCs disappears for increasing ketamine values (Fig. 10A-C). At large enough Ket, the slower LCs does not arise from the branch of faster LCs anymore but instead through a newly emerged Hopf bifurcation at lower values of *I*^Ext^, e.g. for Ket = 0.2 at *I*^Ext^ ≈ −0.0382 ms^−1^ (Fig. 10C). Additionally, the stable part of the *θ*-band LC branch progressively diminishes.

To investigate why the *α*-band power decreases but does not disappear despite the associated LC family vanishing, we looked at the dynamics of an isolated JR model at Ket = 0.3 and different but constant *I*^Ext^ in more detail, as shown in Fig. 11. We selected three values of *I*^Ext^ such that the system resides within the parameter space where a stable focus (*I*^Ext^ = 0.13 ms^−1^ and *I*^Ext^ = 0.3 ms^−1^) or a stable LC (*I*^Ext^ = 0.2 ms^−1^) emerges, as indicated by the bifurcation diagrams in (see Fig. 11A). In the absence of noise (Fig. 11A1-B1), the system residing at the focus does not generate relevant *α*-band oscillations. Instead the PSD is characterized by a low power peak due to the asymptotic spiraling towards the focus. Introducing noise (Fig. 11A2-B2) however induces strong *α*-band oscillations of a magnitude comparable to those observed in the BNM. This holds true independent of the system being on the focus or LC. Moreover in presence of noise, the highest *α*-band powers are observed when *I*^Ext^ is located within the LC regime, whereas perturbations around the foci result in lower power.

**Fig 11.**
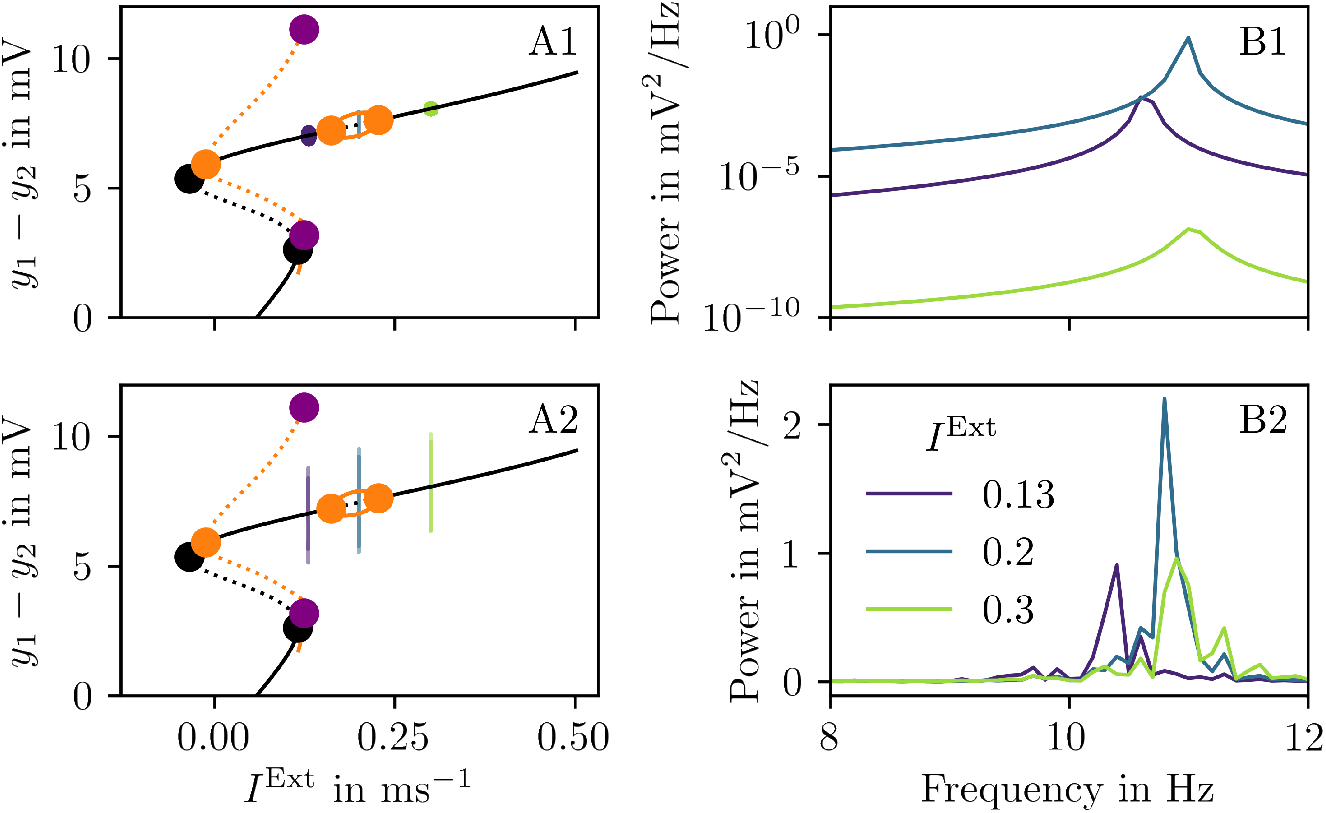
Comparison of different external inputs *I*^Ext^ in one JR node. (**A**) shows the PSP vs. *I*^Ext^ bifurcation diagrams for Ket = 0.3. Three different values of *I*_Ext_ are marked for *I*_Ext_ = 0.13 ms^−1^, *I*_Ext_ = 0.2 ms^−1^ and *I*_Ext_ = 0.3 ms^−1^. The PSP timeseries (*y*_1_ − *y*_2_) is simulated without noise (**A1**) and (**B1**) and with noise in (**A2**) and (**B2**) with noise *σ* = 10^−8^. The PSP values are are superimposed onto the bifurcation diagrams in (**A**) at the corresponding *I*_Ext_. (**B**) show the corresponding frequency vs. power relationship for the three *I*^Ext^ values. Colors and symbols as in Fig. 2.

In short, as the ketamine parameter increases, the region supporting sustained *α*-band oscillations diminishes while the remaining foci become stable 10D. In the full BNM, this causes the *α*-band power to decrease due to the disappearance of the LC. Nevertheless, as shown in Fig. 9, we still observe a non-negligible *α*-band power (Fig. 10E-F) likely due to spiraling fluctuations around the remaining stable foci.

While variations of the LCP naturally change the dynamics at the single-node level, emergent effects on network level need to be studied as well. In the bifurcation diagrams described in Fig. 10, single-node FP and LC branches are represented in (PSP, *I*^Ext^)-space, given constant *I*^Ext^. However, in the BNM interactions between regions enter in *I*^Ext^ such that each region *k* receives distinct time-dependent input 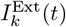 that can lead to more complex behavior of the individual PSP_*k*_(*t*). We can nevertheless use the single JR bifurcation diagram as an estimation of the dynamics of individual nodes nested in the BNM as follows. One can expect regions with larger node strength, i.e., the row sum of the empirical SC, to have a larger 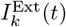 on average. Hence, to capture the variability of dynamics across the BNM, we color-code the time-dependent (PSP_*k*_(*t*), 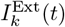 BNM trajectories based on their node strength and superimpose them in the single node bifurcation diagrams. We do not expect the BNM trajectories to precisely follow the single-node attractors but rather remain in their vicinity. This allows us to investigate how the network’s behavior emerges based on the presence or absence of specific single-node attractors. In Fig. 12 we depict this analysis alongside the PSP time series and corresponding PSDs for varying values of the ketamine dose Ket.

**Fig 12.**
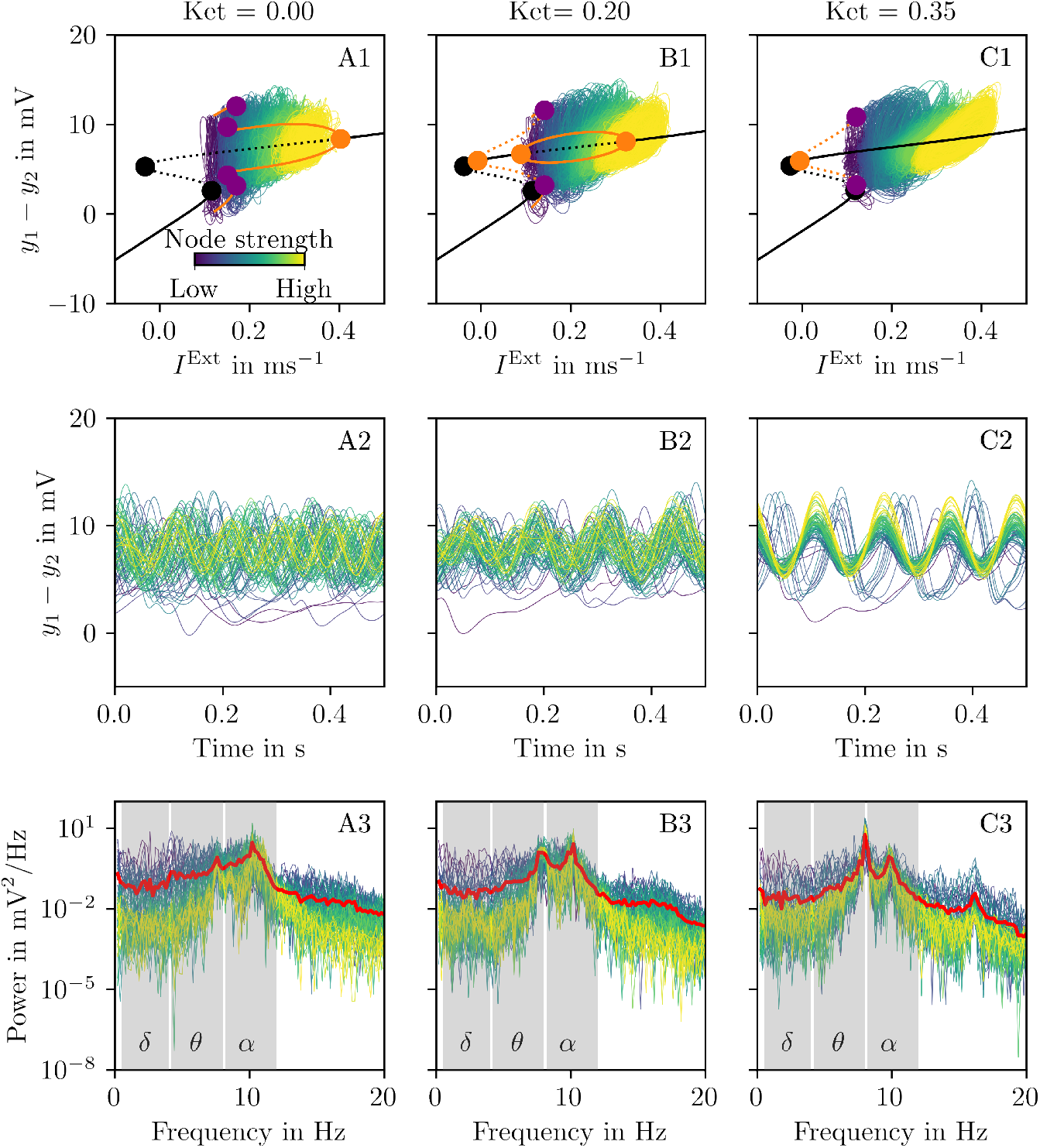
Simulated neural signal at different ketamine doses. Bifurcation diagrams, PSP time series and power spectral density (PSD) are shown for different ketamine doses: no ketamine (Ket = 0.0, **A**), subanesthetic (Ket = 0.20, **B**) and anesthetic (Ket = 0.35, **C**). (**A1-C1**) show the bifurcation diagrams for a single JR model with the respective LCP fixed by the Ket value. For the corresponding network simulations PSP_*k*_ = *y*_1,*k*_ − *y*_2,*k*_ vs. 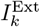 timeseries are obtained for each region *k* and superimposed onto the bifurcation diagram. The color-code depicts the node strength of each region. With increasing Ket the bifurcation types are changing and the *α* LC is diminishing and is replaced by a previously unstable focus that becomes stable. The *θ* LC also progressively diminishes. (**A2-C2**) show the PSP_*k*_(*t*) for each region *k* colored according to their node strength. The PSD computed from PSP_*k*_(*t*) (**A3-C3**) is shown for all regions *k* colored according to node strength and the mean PSD is colored in red.

Indeed, in the absence of Ket (Ket = 0.0, Fig. 12A) we observe that nodes with low node strengths traverse lower *I*^Ext^(*t*) values as compared to nodes with high node strengths (see Fig. 12A2). Hence, poorly connected nodes and well-connected nodes can be associated with *θ*-band and *α*-band oscillations respectively, due to the presence of the corresponding attractors in the single node dynamics. This is further evident from the timeseries in Fig. 12A2 as well as the PSDs depicted in Fig. 12A3 showing an *α*-band peak driven by high strength and a smaller *θ*-band peak driven by low strength nodes, respectively.

For larger Ket values (Ket = 0.20 and Ket = 0.35) we find a decrease in *α*-power (Fig. 12B3 and C3), consistent with the results of the single node analysis in Fig. 11 for which the *α*-power decreases with decreasing size of the region that supports the *α*-band LC. In anesthetic doses the BNM still exhibits non-negligible *α*-band oscillations, stemming from the single node foci together with noise as well as time-varying 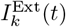. Notably, the BNM synchronizes to some extent with increasing ketamine dose Ket, as can be seen in Fig. 12C2.

## Discussion

In this work, we develop a BNM using the Jansen-Rit neural mass model to explore the effects of ketamine on the brain, specifically in the subcellular and oscillatory domain. Ketamine effects are implemented by modifying the local coupling between subpopulations of neurons, simulating its primary mechanism of action, the NMDAR antagonism. This allows us to capture the synaptic and oscillatory changes observed across different doses of ketamine. Our simulations demonstrate that distinct ketamine-induced NMDARergic alterations on different types of neurons lead to changes in pyramidal cell firing and network oscillations. Specifically, we observed suppressed *α*-oscillations in both sub- and anesthetic regimes and increased *θ*-oscillations in the anesthetic regime consistent with experimental findings in electrophysiological studies [5–7].

There is growing interest in understanding ketamine’s rapid antidepressant effects to develop novel treatments and deepen our understanding of depression. While its molecular targets are relatively well-characterized, the mechanisms by which these targets give rise to higher-order effects remain elusive. While low doses of ketamine show antidepressant and dissociative effects, high doses are used to induce anesthesia. By embedding these dose-dependent effects within one biologically plausible whole-brain framework, we offer a methodology for implementing and testing mechanistic theories. To our knowledge, this is the first whole-brain model depicting ketamine’s NMDAR antagonism across varying doses.

Ketamine research spans various species, each offering distinct advantages and constraints based on the achievable invasiveness and resolution. Since ketamine directly interacts with the NMDAR, rodent studies can provide insights into its impact on neurotransmission by measuring glutamate concentration [33]. Conversely, in humans glutamate concentration can only be inferred indirectly via magnetic resonance spectroscopy. Furthermore, rodent studies provide microscale insights while human studies are restricted to the meso- or macroscale via e.g. EEG or fMRI. The leveraging of cross-species information to link microscale mechanisms to meso-and large-scale phenomena is complicated. The differences in ketamine doses further complicates this translation. Whole-brain computational models can address this challenge by embedding rodent-derived mechanisms into human-scale frameworks, enabling robust theory testing and validation against empirical human data. In this study, we implement the disinhibition theory of ketamine, which suggests that low doses preferentially target IINs, reducing their inhibition and thereby disinhibiting pyramidal cells [13]. We also implemented a simpler model of uniform NMDAR antagonism to compare it with. Our results demonstrate that only the disinhibition model aligns with the effects found in literature. Using a whole-brain model, we abstracted receptor-level effects to mesoscale population dynamics, modeling ketamine’s impact as disrupted local excitatory and inhibitory coupling. By applying sigmoidal transformations to represent receptor antagonism, we uncovered how localized disinhibition propagates to global network-level changes. Furthermore, we systematically explored the local dynamics, providing insights into how ketamine alters brain-wide activity.

The combination of computational modeling with in-depth bifurcation analysis in pharmacological modeling provides mechanistic insights into how changes in parameters impact activity on a network level. It identifies critical points where changes in local coupling parameters cause shifts in brain dynamics, such as from stable fixed points to oscillatory regimes, aiding in understanding ketamine-induced state transitions. It also gives the opportunity to have more predictive power for testing other pharmacological interventions by taking the underlying local dynamics into account showing the possibilities of the model. This provides a framework to predict potential brain responses to other pharmacological interventions or changes in the model. Nevertheless, there are network effects that are not predictable by looking at the local dynamics. Furthermore, cellular-level pharmacological actions can be linked to macroscopic phenomena like altered EEG rhythms offering a biologically grounded explanation of drug effects. Moreover, bifurcation analysis highlights which parameter ranges are most sensitive to ketamine’s effects, helping to refine the model. The biological interpretation of bifurcation analysis in the context of ketamine’s effect on the brain lies in its ability to connect mathematical dynamics with observable neural phenomena. By systematically analyzing how changes in local coupling parameters between subpopulations (mimicking ketamine’s NMDAR-antagonism) lead to shifts in brain states, bifurcation analysis can elucidate mechanisms underlying altered oscillatory activity, such as disrupted alpha rhythms. These dynamics correlate directly with ketamine-induced cognitive effects, providing a mechanistic understanding of its impact on neural coordination and brain network functionality.

We present a flexible model that captures ketamine’s effects on NMDAR-mediated coupling across varying doses. However, there are several limitations to this study. One key limitation is that our model is restricted to generating oscillatory activity in the *α*- and *θ*-bands, omitting changes in *γ* activity that have been consistently observed in empirical studies [34, 35]. To improve biological plausibility, future work could address these multifrequency dynamics by coupling the Jansen-Rit model with a PING model, which is capable of producing *γ* oscillations [36]. Another limitation lies in the behavior of the Jansen-Rit model under low local coupling conditions, where node time series become increasingly synchronized. In contrast, empirical studies show both synchronization and desynchronization in EEG signals depending on frequency bands and brain regions (e.g. *α* desynchronization and *θ* synchronization [7]). This problem might arise from our sole focus on local implementations of ketamine’s effects. Future refinements could explore the inclusion of global effects, such as ketamine-induced reductions in long-range coupling, to better capture these dynamics. Our model focuses on ketamine’s primary mechanism of action which is the NMDAR antagonism but ketamine is a dirty drug that acts on multiple receptor systems and produces active metabolites [37, 38]. This complexity suggests that not all observed effects can be attributed solely to NMDAR antagonism. Future studies could incorporate alternative theories of ketamine’s mechanisms of action [39] and the roles of its metabolites [40], enabling differential analyses to better understand its multifaceted effects. Future investigations could also test the generalizability of the sigmoidal dose dependencies observed here across other computational frameworks. Additionally, the model could be extended to study other NMDAR-modulating therapeutics, such as memantine or diseases targeting the NDMAR, such as antibody-mediated NMDAR encephalitis. This enables comparisons of NMDAR-blocking efficiencies and behavioral outcomes. Validation through experimental electrophysiology remains critical to refining the model. By incorporating personalized structural connectomes, this framework holds potential for patient-specific modeling, offering insights into individual responsiveness to ketamine treatment. Ultimately, this approach could contribute to precision medicine, identifying patients most likely to benefit from ketamine therapy.

Our model helps bridge the gap between micro- to meso-and large-scale effects of ketamine, demonstrating significant potential in predicting its effects and guiding future studies of other pharmacological agents. This work has broad clinical implications, particularly as ketamine is under investigation as treatment for a range of neuropsychiatric disorders, including bipolar disorder, anxiety spectrum disorders, and post-traumatic stress disorder [41–43]. Our findings highlight the utility of computational modeling as a framework for testing and validating proposed pharmacological mechanisms of action. By leveraging insights from rodent to human studies, this approach provides a pathway to understanding how micro-level effects propagate to higher-order systems. This knowledge can advance the search for pharmacological treatments for the increasing burden of neuropsychiatric disorders while also enhancing our understanding of the disorders themselves.

## Acknowledgments

PR acknowledges support by EU Horizon Europe program Horizon BRIDGE (101219311), EBRAINS2.0 (101147319), Virtual Brain Twin (101137289), EBRAINS-PREP 101079717, AISN 101057655, EBRAIN-Health 101058516, EIC grant PHRASE 101058240, by the Digital Europe Programme TEF-Health (101100700), SHAIPED (101195135), CoordinaTEF (101168074) German Research Foundation SFB 1436 (project ID 425899996); SFB 1315 (project ID 327654276); SFB 936 (project ID 178316478; SFB-TRR 295 (project ID 424778381); SPP Computational Connectomics RI 2073/6-1, RI 2073/10-2, RI 2073/9-1; DFG Clinical Research Group BECAUSE-Y 504745852, Berlin University Alliance OpenMake, the Virtual Research Environment at the Charité Berlin and EBRAINS Health Data Cloud and the Berlin Institute of Health and Foundation Charité. JM acknowledges support by the Deutsche Forschungsgemeinschaft (DFG, German Research Foundation) - Project-ID 424778381 - TRR 295. We acknowledge the assistance of ChatGPT, an AI language model developed by OpenAI, which was used solely for language refinement and proofreading purposes. All scientific content and interpretations remain the sole responsibility of the authors.

## References

1. Alnefeesi Y, Chen-Li D, Krane E, Jawad Y, Rodrigues N, Ceban F, et al. Real-world effectiveness of ketamine in treatment-resistant depression: A systematic review meta-analysis. Journal of Psychiatric Research. 2022;151. doi:10.1016/j.jpsychires.2022.04.037.

2. Dehestani S, Mohammadpour A, Sadjadi SA, Sathyapalan T, Sahebkar A. Clinical use of ketamine in psychiatric disorders. Annales Médico-psychologiques, revue psychiatrique. 2022;181. doi:10.1016/j.amp.2022.05.008.

3. Wellington N, Bouças A, Lagopoulos J, Quigley B, Kuballa A. Molecular pathways of ketamine: A systematic review of immediate and sustained effects on PTSD. Psychopharmacology. 2025;242:1197–1243. doi:10.1007/s00213-025-06756-4.

4. Moghaddam B, Adams B, Verma A, Daly D. Activation of Glutamatergic Neurotransmission by Ketamine: A Novel Step in the Pathway from NMDA Receptor Blockade to Dopaminergic and Cognitive Disruptions Associated with the Prefrontal Cortex. The Journal of neuroscience : the official journal of the Society for Neuroscience. 1997;17:2921–7. doi:10.1523/JNEUROSCI.17-08-02921.1997.

5. Zacharias N, Müller F, Lammers-Lietz F, Saleh A, London M, Boer P, et al. Ketamine effects on default mode network activity and vigilance: A randomized, placebo-controlled crossover simultaneous fMRI/EEG study. Human Brain Mapping. 2019;41. doi:10.1002/hbm.24791.

6. Akeju O, Song A, Hamilos A, Pavone K, Flores F, Brown E, et al. Electroencephalogram Signatures of Ketamine-Induced Unconsciousness. Clinical Neurophysiology. 2016;127. doi:10.1016/j.clinph.2016.03.005.

7. Vlisides P, Bel-Bahar T, Lee U, Li D, Kim H, Janke E, et al. Neurophysiologic Correlates of Ketamine Sedation and Anesthesia: A High-density Electroencephalography Study in Healthy Volunteers. Anesthesiology. 2017;127. doi:10.1097/ALN.0000000000001671.

8. Tian F, Lewis L, Zhou D, Balanza G, Paulk A, Zelmann R, et al. Characterizing brain dynamics during ketamine-induced dissociation and subsequent interactions with propofol using human intracranial neurophysiology. Nature Communications. 2023;14. doi:10.1038/s41467-023-37463-3.

9. Antkowiak B. Different Actions of General Anesthetics on the Firing Patterns of Neocortical Neurons Mediated by the GABAAReceptor. Anesthesiology. 1999;91(2):500–511. doi:10.1097/00000542-199908000-00025.

10. Homayoun H, Moghaddam B. NMDA Receptor Hypofunction Produces Opposite Effects on Prefrontal Cortex Interneurons and Pyramidal Neurons. Journal of Neuroscience. 2007;27(43):11496–11500. doi:10.1523/JNEUROSCI.2213-07.2007.

11. Zhang B, Yang X, Ye L, Liu R, Ye B, Du W, et al. Ketamine activated glutamatergic neurotransmission by GABAergic disinhibition in the medial prefrontal cortex. Neuropharmacology. 2021;194:108382. doi:10.1016/j.neuropharm.2020.108382.

12. Abdallah C, De Feyter H, Averill L, Jiang L, Averill C, Chowdhury GMI, et al. The effects of ketamine on prefrontal glutamate neurotransmission in healthy and depressed subjects. Neuropsychopharmacology. 2018;43. doi:10.1038/s41386-018-0136-3.

13. Kohtala S. Ketamine—50 years in use: from anesthesia to rapid antidepressant effects and neurobiological mechanisms. Pharmacological Reports. 2021;73. doi:10.1007/s43440-021-00232-4.

14. Cichon J, Wasilczuk A, Looger L, Contreras D, Kelz M, Proekt A. Ketamine triggers a switch in excitatory neuronal activity across neocortex. Nat Neurosci. 2023;26(1):39–52. doi:10.1038/s41593-022-01203-5.

15. Izumi Y, Zorumski C. Metaplastic Effects of Subanesthetic Ketamine on CA1 Hippocampal Function. Neuropharmacology. 2014;86. doi:10.1016/j.neuropharm.2014.08.002.

16. Rao S, Williams G, Goldman-Rakic P. Correlated Discharges Among Putative Pyramidal Neurons and Interneurons in the Primate Prefrontal Cortex. Journal of neurophysiology. 1999;81:1903–1916. doi:10.1152/jn.1999.81.4.1903.

17. Constantinidis C, Goldman-Rakic P. Correlated Discharges Among Putative Pyramidal Neurons and Interneurons in the Primate Prefrontal Cortex. Journal of neurophysiology. 2003;88:3487–97. doi:10.1152/jn.00188.2002.

18. Gerhard D, Pothula S, Liu RJ, Wu M, Li XY, Girgenti M, et al. GABA interneurons are the cellular trigger for ketamine’s rapid antidepressant actions. Journal of Clinical Investigation. 2019;130. doi:10.1172/JCI130808.

19. Krystal J, D’Souza D, Mathalon D, Perry E, Belger A, Hoffman R. Krystal JH, D’Souza DC, Mathalon D, Perry E, Belger A, Hoffman R. NMDA receptor antagonist effects, cortical glutamatergic function, and schizophrenia: toward a paradigm shift in medication development. Psychopharmacology (Berl) 169: 215-233. Psychopharmacology. 2003;169:215–33. doi:10.1007/s00213-003-1582-z.

20. Kotermanski SE, Johnson JW. Mg2+ imparts NMDA receptor subtype selectivity to the Alzheimer’s drug memantine. the official journal of the Society for Neuroscience. 2009;29:2774–2779. doi:10.1523/JNEUROSCI.3703-08.2009.

21. Widman A, Mcmahon L. Disinhibition of CA1 pyramidal cells by low-dose ketamine and other antagonists with rapid antidepressant efficacy. Proceedings of the National Academy of Sciences. 2018;115:201718883. doi:10.1073/pnas.1718883115.

22. Ali F, Gerhard D, Sweasy K, Pothula S, Pittenger C, Duman R, et al. Ketamine disinhibits dendrites and enhances calcium signals in prefrontal dendritic spines. Nature Communications. 2020;11:1234567890. doi:10.1038/s41467-019-13809-8.

23. Sacha M, Tesler F, Cofre R, Destexhe A. A computational approach to evaluate how molecular mechanisms impact large-scale brain activity. Nature Computational Science. 2025;5:405–417. doi:10.1038/s43588-025-00796-8.

24. Adam E, Kowalski M, Akeju O, Miller E, Brown E, Mccarthy M, et al. Ketamine can produce oscillatory dynamics by engaging mechanisms dependent on the kinetics of NMDA receptors. Proceedings of the National Academy of Sciences of the United States of America. 2024;121:e2402732121. doi:10.1073/pnas.2402732121.

25. Shaw A, Muthukumaraswamy S, Saxena N, Sumner R, Adams N, Moran R, et al. Generative modelling of the thalamo-cortical circuit mechanisms underlying the neurophysiological effects of ketamine. NeuroImage. 2020;221:117189. doi:10.1016/j.neuroimage.2020.117189.

26. Jansen BH, Rit VG. Electroencephalogram and visual evoked potential generation in a mathematical model of coupled cortical columns. Biological cybernetics. 2005;73:357–366. doi:10.1007/BF00199471.

27. Van Essen D, Smith S, Barch D, Behrens T, Yacoub E, Ugurbil K. The WU-Minn Human Connectome Project: an overview. NeuroImage. 2013;80. doi:10.1016/j.neuroimage.2013.05.041.

28. Glasser M, Sotiropoulos S, Wilson J, Coalson T, Fischl B, Andersson J, et al. The Minimal Preprocessing Pipelines for the Human Connectome Project. NeuroImage. 2013;80:105. doi:10.1016/j.neuroimage.2013.04.127.

29. Desikan R, Ségonne F, Fischl B, Quinn B, Dickerson B, Blacker D, et al. An automated labeling system for subdiving the human cerebral cortex on MRI scans into gyral based regions of interest. NeuroImage. 2006;31:968–80. doi:10.1016/j.neuroimage.2006.01.021.

30. Fischl B, Kouwe A, Destrieux C, Ségonne F, Salat D, Busa E, et al. Automatically Parcellating the Human Cerebral Cortex. Cerebral cortex (New York, NY : 1991). 2004;14:11–22.

31. Sanz-Leon P, Knock S, Woodman M, Domide L, Mersmann J, McIntosh A, et al. The Virtual Brain: a simulator of primate brai network dynamics. Frontiers in neuroinformatics. 2013;7:10. doi:10.3389/fninf.2013.00010.

32. Doedel E, Fairgrieve T, Sandstede B, Champneys A, Kuznetsov Y, Wang X. AUTO-07P: Continuation and bifurcation software for ordinary differential equations. 2007;.

33. Cifuentes-Castro V, Valenzuela C, Sanchez J, Pardo-Peña K, López-Pérez S, Ibarra J, et al. An Update of the Classical and Novel Methods Used for Measuring Fast Neurotransmitters During Normal and Brain Altered Function. Current Neuropharmacology. 2014;12. doi:10.2174/1570159X13666141223223657.

34. Shaw A, Saxena N, Jackson L, Hall J, Singh K, Muthukumaraswamy S. Ketamine amplifies induced gamma frequency oscillations in the Human cerebral cortex. European Neuropsychopharmacology. 2015;25. doi:10.1016/j.euroneuro.2015.04.012.

35. Curic S, Andreou C, Steinmann S, Thiebes S, Polomac N, Haaf M, et al. Ketamine Alters Functional Gamma and Theta Resting-State Connectivity in Healthy Humans: Implications for Schizophrenia Treatment Targeting the Glutamate System. Frontiers in Psychiatry. 2021;12:671007. doi:10.3389/fpsyt.2021.671007.

36. Sanchez-Todo R, Bastos A, Sola E, Mercadal B, Santarnecchi E, Miller E, et al. A physical neural mass model framework for the analysis of oscillatory generators from laminar electrophysiological recordings. NeuroImage. 2023;270:119938. doi:10.1016/j.neuroimage.2023.119938.

37. Zanos P, Gould T. Mechanisms of ketamine action as an antidepressant. Molecular Psychiatry. 2018;23. doi:10.1038/mp.2017.255.

38. Sial O, Parise E, Parise L, Gnecco T, Bolaños C. Ketamine: The final frontier or another depressing end? Behavioural Brain Research. 2020;383:112508. doi:10.1016/j.bbr.2020.112508.

39. Miller O, Moran J. Two Cellular Hypotheses Explaining Ketamine’s Antidepressant Actions: Direct Inhibition and Disinhibition. Neuropharmacology. 2016;100. doi:10.1016/j.neuropharm.2015.07.028.

40. Zanos P, Moaddel R, Morris P, Georgiou P, Fischell J, Elmer G, et al. NMDAR inhibition-independent antidepressant actions of ketamine metabolites. Nature. 2016;533. doi:10.1038/nature17998.

41. Bahji A, Zarate C, Vazquez G. Ketamine for Bipolar Depression: A Systematic Review. The international journal of neuropsychopharmacology. 2021;24. doi:10.1093/ijnp/pyab023.

42. Whittaker E, Dadabayev A, Joshi S, Glue P. Systematic review and meta-analysis of randomized controlled trials of ketamine in the treatment of refractory anxiety spectrum disorders. Therapeutic Advances in Psychopharmacology. 2021;11:204512532110567. doi:10.1177/20451253211056743.

43. Jumaili W, Trivedi C, Chao T, Kubosumi A, Jain S. The safety and efficacy of Ketamine NMDA receptor blocker as a therapeutic intervention for PTSD review of a randomized clinical trial. Behavioural Brain Research. 2022;424:113804. doi:10.1016/j.bbr.2022.113804.

